# The U-method: Leveraging expression probability for robust biological marker detection

**DOI:** 10.64898/2026.03.31.715470

**Authors:** Y. Stein, H. Lavon, M. Hindi Malowany, L. Arpinati, R. Scherz-Shouval

## Abstract

Reliable identification of cluster-defining markers is fundamental to single-cell transcriptomic analysis, yet current approaches often rely on average expression differences, which can dilute biologically informative signals in sparse and heterogeneous data. Here we introduce the U-method, a fast probability-based framework for identifying uniquely expressed genes (UEGs) by contrasting a gene’s expression probability within a cluster with its highest expression probability in any other cluster. This highest-probability comparison prioritizes detection consistency over expression magnitude, resulting in markers that consistently identify cell populations across independent datasets analyzed at comparable clustering resolutions. Applied to colorectal, breast, pancreatic, and lung cancer single-cell RNA-sequencing datasets, the U-method identifies canonical lineage markers together with additional genes showing clear cluster specificity. When projected onto Visium HD spatial transcriptomics data using only raw average expression of top UEGs, these signatures reveal coherent and biologically interpretable tissue organization without the need for smoothing, deconvolution, or model-based spatial inference. These results position the U-method as a practical implementation of detection consistency, enabling robust marker discovery and spatial interpretation in single-cell analysis.

## Introduction

Single-cell RNA sequencing (scRNA-seq) enables comprehensive measurement of gene expression across diverse cell types, offering a detailed molecular view of tissue composition^1–7^. When combined with spatial transcriptomics, these approaches provide a powerful means to map cellular identity, function, and organization within intact tissues^2,8–12^. To fully exploit this potential for distinguishing cell populations and decoding tissue complexity, robust marker genes are essential^13,14^. However, high-dimensional inference in such settings is often highly sensitive to modeling assumptions and analytical choices, limiting the stability and reproducibility of resulting conclusions^15^. This sensitivity is particularly consequential in differential expression (DE) analysis.

Despite major advances in profiling technologies, differential expression (DE) analysis often remains a bottleneck in the interpretation pipeline ^13,14^. Most DE analysis methods rely on average expression to define cluster identity, capturing expression magnitude but not how consistently a gene marks a population across individual cells. This emphasis can blur biologically meaningful distinctions between closely related populations^13,16–19^. Downstream analyses, from cell type annotation to spatial deconvolution, depend on clean cluster boundaries and uniquely expressed markers. When these are missing or misidentified, even subtle shifts in expression can lead to incorrect biological interpretation^14,15,19^.

While normalization can reduce some technical variation, the heterogeneity and sparsity of scRNA-seq data still pose major challenges for marker detection^7,13,16,20^. A wide range of approaches, including normalization, batch correction, and imputation, attempt to stabilize the expression matrix, but do not explicitly address whether candidate markers are consistently detected across cells within a population^4,7^. This issue becomes even more pronounced in newer spatial platforms such as 10x Genomics Visium HD, where higher resolution increases the need for consistent and reproducible markers across samples and conditions^10,21,22^.

Many standard differential-expression frameworks assume continuous, well-behaved expression distributions. However, single-cell data are sparse and zero-inflated, with many biologically informative genes effectively “on” in one population and absent in others^13,20,23^. While existing methods can capture gradients and trajectories, they are highly sensitive to normalization choices and read-count variation, which can obscure crisp, cluster-defining signals^13,14^. A probability-based perspective addresses this limitation by prioritizing a complementary property of gene expression: detection consistency across cells. By quantifying how often a gene is expressed in a given cluster relative to others, detection likelihood provides a stable and interpretable basis for identifying markers that reliably distinguish cell populations.

Here, we present the U-method, a fast and interpretable framework for identifying cluster-specific markers and projecting them into spatial context based on their expression probability, without requiring normalization or scaling. The U-method ranks genes by differences in detection probability across predefined clusters, prioritizing markers that are consistently expressed within a population relative to others. Rather than replacing magnitude-based approaches, it provides a complementary axis for marker identification that emphasizes detection consistency. Applied across multiple publicly available cancer datasets and projected onto Visium HD datasets, U-method derived markers support robust cell-type and subpopulation annotation and enable coherent spatial projection. Together, these results highlight detection consistency as a general principle for marker discovery, with the U-method providing a practical and scalable implementation.

## Results

### The U-method identifies consistent, biologically interpretable marker genes in scRNA-seq data

A central goal of marker identification in single-cell transcriptomics is to identify genes that reliably distinguish cell populations. In sparse single-cell data, expression magnitude alone is often unstable, as it can be driven by a small number of high-count cells. To address this, the U-method ranks markers based on detection consistency.

The U-method is applied to a labeled single-cell dataset and evaluates each gene at the cluster level by contrasting its expression probability within a given population with the highest expression probability observed in any other population. This comparison yields a ranked list of cluster-specific markers, which can be carried forward into downstream analyses, including spatial projection (Figure 1A).

**Figure 1.**
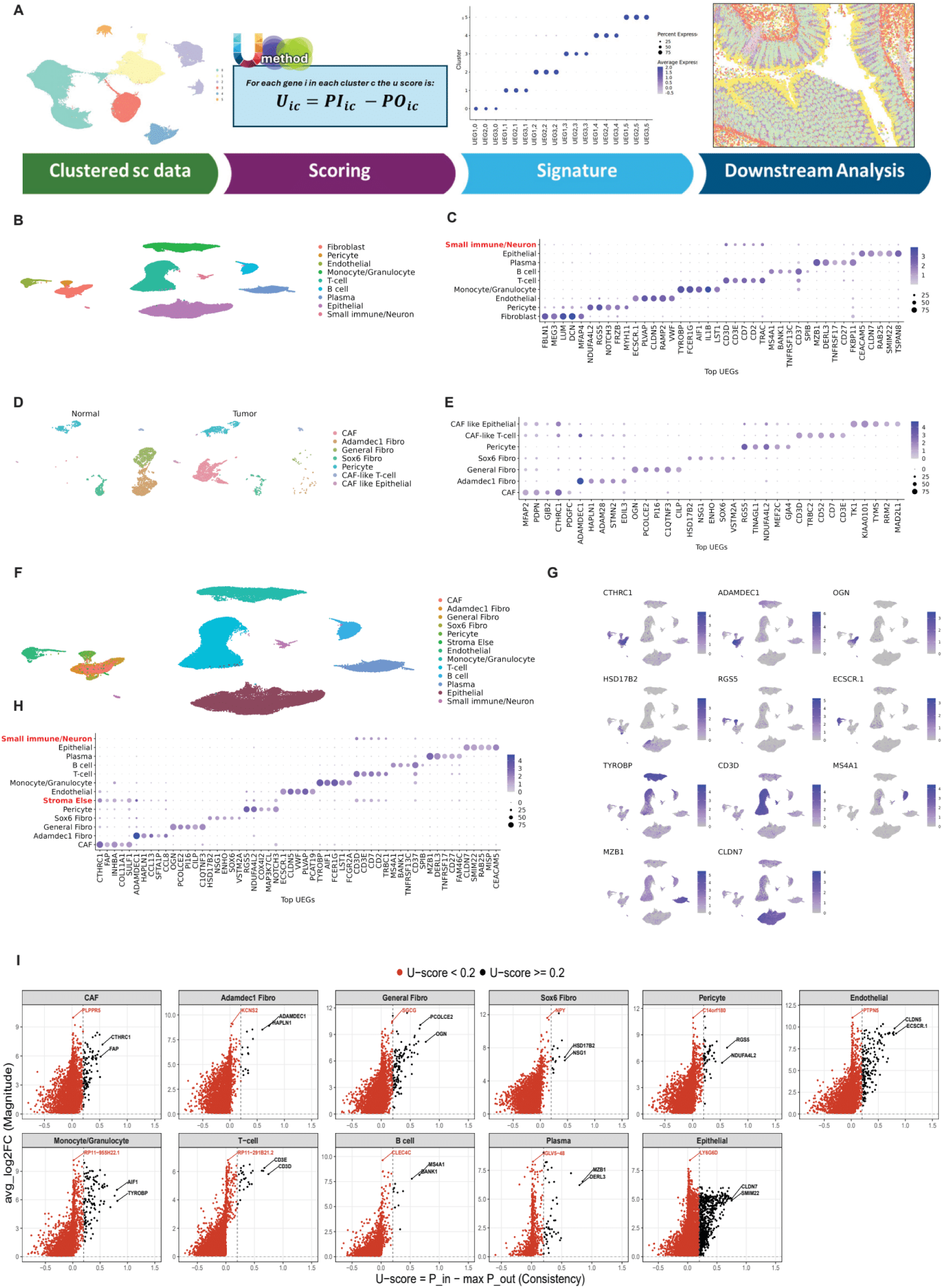
The U-method identifies uniquely expressed genes (UEGs) characterizing cell subsets in scRNA-seq data from colorectal cancer. (A) Schematic representation of the U-method workflow. Cluster-specific marker genes are identified from scRNA-seq data using a probability-based scoring function (P_in_ − P_out_). These markers can be used for downstream analyses, including spatial projection. (B) UMAP projections of scRNA-seq data from human colorectal cancer (CRC) and adjacent normal colon tissue^2^, analyzed by Seurat and annotated by major cell type. (C) Unscaled Dot plot of the top uniquely expressed genes (UEGs) per cluster. Dot size reflects the proportion of cells expressing each gene. (D) UMAP projections of scRNA-seq data of stromal cells, divided into cells from tumor and from normal adjacent colon tissue annotated by stromal subcluster. (E) Unscaled Dot plot of the top UEGs in each stromal cluster. (F) UMAP projections of the colon scRNA-seq dataset^2^ showing stromal subclusters on the general all-cell UMAP. (G-H) Unscaled Dot plot and Feature plots of the top UEGs per cluster. (I) Scatter plots showing U-scores (x-axis) and Log2 fold change values (y-axis; computed from differential expression analysis) per gene per cluster. Genes with U-scores greater than 0.2 are highlighted.

To demonstrate the performance of the U-method, we analyzed a colorectal cancer (CRC) scRNA-seq dataset comprising 23 tumors and 10 matched normal mucosa samples^2^. Using Seurat clustering^24,25^, we identified nine major cell populations which were visualized by UMAP embedding (Figure 1B). Applying the U-method to these clusters identified robust canonical lineage markers, enabling the classification of clusters to known cell types including epithelial, fibroblast, endothelial, and immune lineages (Figure 1C, Supplementary Table 1): CD3D for T cells, MS4A1 for B cells, MZB1 for plasma cells, and TYROBP for myeloid cells. Dot plots of these markers showed strong diagonal enrichment, with each marker expressed predominantly in its cluster of origin (Figure 1C). By recovering lineage-defining genes directly from observed expression probabilities without predefined marker sets, supervised labels, or latent-space models, the U-method provides a route to cell-type annotation. Moreover, it detects additional genes with more subtle yet consistently enriched expression, broadening the catalog of markers available for downstream analyses.

To test the robustness of the U-method, we applied it to a subset of the data - the stromal fraction, focusing on fibroblasts and pericytes. These stromal cells were reanalyzed using the same Seurat pipeline, and the U-method was then applied to the resulting subclusters (Figure 1D). The analysis recovered five well-described stromal populations in the colon, with high-scoring U-markers corresponding to established cell-defining genes^2,8^. For example, OGN and PI16 marked general fibroblasts while MFAP2 and PDPN identified cancer-associated fibroblasts (CAFs). ADAMDEC1 defined an ADAMDEC1⁺ fibroblast population, SOX6 labeled upper-crypt fibroblasts, and RGS5 together with NOTCH3 identified pericytes (Figure 1E). We also identified two small mixed groups, which we annotated CAF-like T cells and CAF-like epithelial cells due to their expression of a mix of broad stromal markers and broad signatures from other non-stromal cells.

Following stromal subclustering, all stromal populations were projected back onto the full dataset, and the two mixed groups were merged into a single “Stroma Else” category (Figure 1F). This decision was motivated by the observation that these clusters exhibited concurrent expression of marker genes associated with multiple distinct stromal and non-stromal cell states, rather than a coherent, cluster-specific expression program. Such mixed profiles likely reflect dividing, transitional, hypoxic, or doublet cells and do not support stable identification of uniquely expressed genes. To prevent these mixed populations from diluting the background probability estimates, they were excluded from the out-of-cluster (P_out_) calculation used for U-score computation, while remaining part of the dataset (Figure 1F-H). This exclusion sharpened the definition of cluster-specific UEGs in the remaining stromal populations, with the excluded clusters retained in the UMAP to reflect their non-specific UEG expression.

Using this refined annotation, the U-method enabled consistent and confident identification of stromal cell states, including CAFs, general fibroblasts, ADAMDEC1⁺ fibroblasts, SOX6⁺ fibroblasts, and pericytes (Figure 1G-H). Marker assignments remained stable, or became more refined in the case of CAFs, after relabeling and reapplying the U-method to the full dataset.

Unlike many differential expression methods used for defining markers, which prioritize expression magnitude, the U-method defines marker specificity based on detection consistency across cells within a cluster. To relate these two perspectives, we compared consistency-based and magnitude-based marker definitions by visualizing gene-level U-scores alongside log2 fold-change values for all genes within each cluster (Figure 1I). Across clusters, many genes with high fold-change values showed U-scores near zero, indicating that large expression differences do not necessarily correspond to consistent detection across cells. A smaller subset of genes exhibited elevated U-scores and was also associated with higher fold-change values, as expected given that presence-absence differences contribute to expression magnitude. Together, these observations indicate that genes with strong log2 fold-change signals and genes with strong U-score signals capture different aspects of marker behavior.

Aggregation-based approaches are commonly used to mitigate the effects of sparsity in single-cell data by stabilizing expression magnitude estimates through summarizing expression across samples. To examine whether such aggregation aligns magnitude-based marker definitions with detection-based specificity, we compared pseudobulk fold changes with U-scores across clusters. Although aggregation reduced variability in magnitude estimates, the distinction between pseudobulk-derived measures and detection-based U-scores persisted (Supplementary Figure 1), indicating that expression magnitude and detection consistency capture separable aspects of marker specificity.

### The U-method identifies stable, identity-defining markers across independent datasets

To evaluate the generalizability and reproducibility of the U-method across independent datasets, we applied it to four publicly available scRNA-seq datasets: two from breast cancer^5,7^ (Figure 2A-D) and two from pancreatic ductal adenocarcinoma (PDAC)^4,6^ (Figure 2E-H). Each dataset was processed using a standard Seurat workflow, and the same U-method parameters were applied without dataset-specific adjustment. For visualization, we displayed the top five uniquely expressed genes per cluster, filtered to include only those genes with U-scores greater than 0.2, mirroring the workflow utilized in the colorectal dataset (Figure 2C-D, 2G-H). Across datasets, the U-method identified robust canonical markers for the different clusters, many of which were shared between the different datasets, suggesting that the U-method consistently identifies the same cluster-specific markers across tumor types.

**Figure 2.**
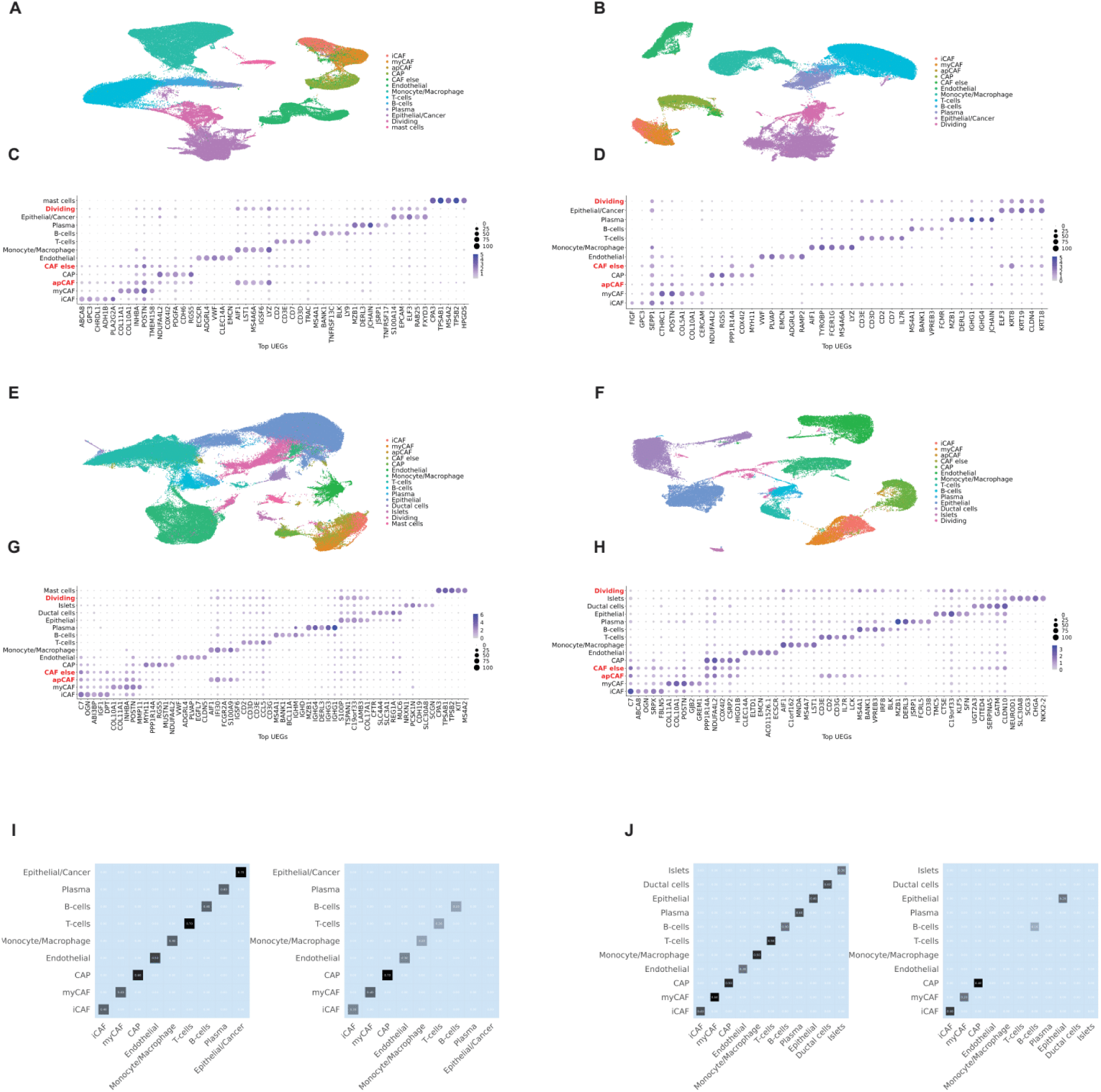
The U-method identifies reproducible markers across breast and pancreatic cancer datasets. (A-B) UMAP representations of two independent scRNA-seq breast cancer patient datasets^5,7^ processed using the same Seurat workflow and annotated by cell type. (C-D) Unscaled Dot plots of the breast cancer datasets (U-score > 0.2). (E-F) UMAP representations of two independent scRNA-seq pancreatic ductal adenocarcinoma (PDAC) patient datasets^4,6^, processed as in (A-B). (G-H) Unscaled Dot plots of the two PDAC datasets showing the top five U-score markers per cluster (U-score > 0.2). (I) Heatmap showing, for each matched cell type, the overlap likelihood of shared genes between the top ten markers derived independently from two breast cancer scRNA-seq datasets. The left heatmap reports overlap based on U-method markers, where genes were ordered by U-score within each cluster and the top ten genes were used to compute overlap. The right heatmap reports overlap based on markers identified using Seurat’s Wilcoxon-based differential expression analysis with |log2 fold change| > 2 and adjusted p-value < 0.05, using the top ten genes per cluster. (J) Heatmap showing overlap likelihood of the top ten markers across matched cell types between two PDAC scRNA-seq datasets, computed using the same U-method and Wilcoxon-based marker selection procedures as in (I).

To quantitively assess cross-dataset reproducibility of consistency-based markers, we quantified the overlap of U-method markers between datasets by comparing the top ten ranked genes per cluster across matched cell types in matched tumor types. Genes were ordered by U-score within each cluster, without applying a U-score threshold (Figure 2I–J, Left panels). The resulting overlap matrices showed a pronounced diagonal structure, with maximal overlap between corresponding clusters across datasets and no overlap outside the diagonal. This pattern indicates stable correspondence of cluster identities and reproducible marker selection when genes are ranked by detection consistency.

We performed the same analysis using markers identified by Seurat’s Wilcoxon rank-sum test applied to the same datasets. Markers were selected using |log2 fold change| > 2 and adjusted p-value < 0.05, and the top-ranked genes per cluster were used to compute overlap (Figure 2I–J, Right panels). In contrast to the U-method, overlap patterns were more variable across datasets, reflecting differences in the genes prioritized by magnitude-based ranking.

A focused analysis in colorectal tissue illustrates the source of this difference (Supplementary Figure 2). Under standard Wilcoxon filtering, several genes with large magnitude differences were detected in only a small fraction of cells within the target cluster (Supplementary Figure 2A) and overlap between the top 10 Wilcoxon markers and the top 10 unfiltered U-markers was low across most clusters (Supplementary Figure 2B). Applying an additional filter requiring expression in more than 25% of cells (pct.1 > 0.25) increased agreement with U-method marker sets (Supplementary Figures 2C-D). This additional step effectively introduces detection frequency as a post hoc criterion, but may still retain genes expressed across multiple clusters, limiting their specificity. In contrast, the U-method incorporates detection consistency directly into marker scoring, providing a unified framework for ranking genes by how reliably they distinguish a cell population.

### The U-method captures spatial organization and highlights biologically meaningful structures

Biologically meaningful clusters are expected to form coherent structures *in-situ*. To evaluate whether U-method markers identified through scRNA-seq analysis inform biologically meaningful clusters in tissue space, we projected the top UEGs from the CRC scRNA-seq dataset onto VisiumHD sections from 3 tumor and 2 normal adjacent colon samples (https://www.10xgenomics.com/platforms/visium/product-family/dataset-human-crc). For each cluster, we computed the raw average expression of its top five U-markers, or all available markers when fewer were present, and plotted these values directly without smoothing, scaling, or deconvolution, preserving raw signals (Figure 3A-B). Across patients, these projections reproduced coherent epithelial, fibroblast, immune, and endothelial compartments, supporting the robustness of the probability-based markers when moved from non-spatial to spatial single-cell data. The U-markers performed as robustly as canonical lineage genes and, in some cases outperformed them (Figure 3C-D).

**Figure 3.**
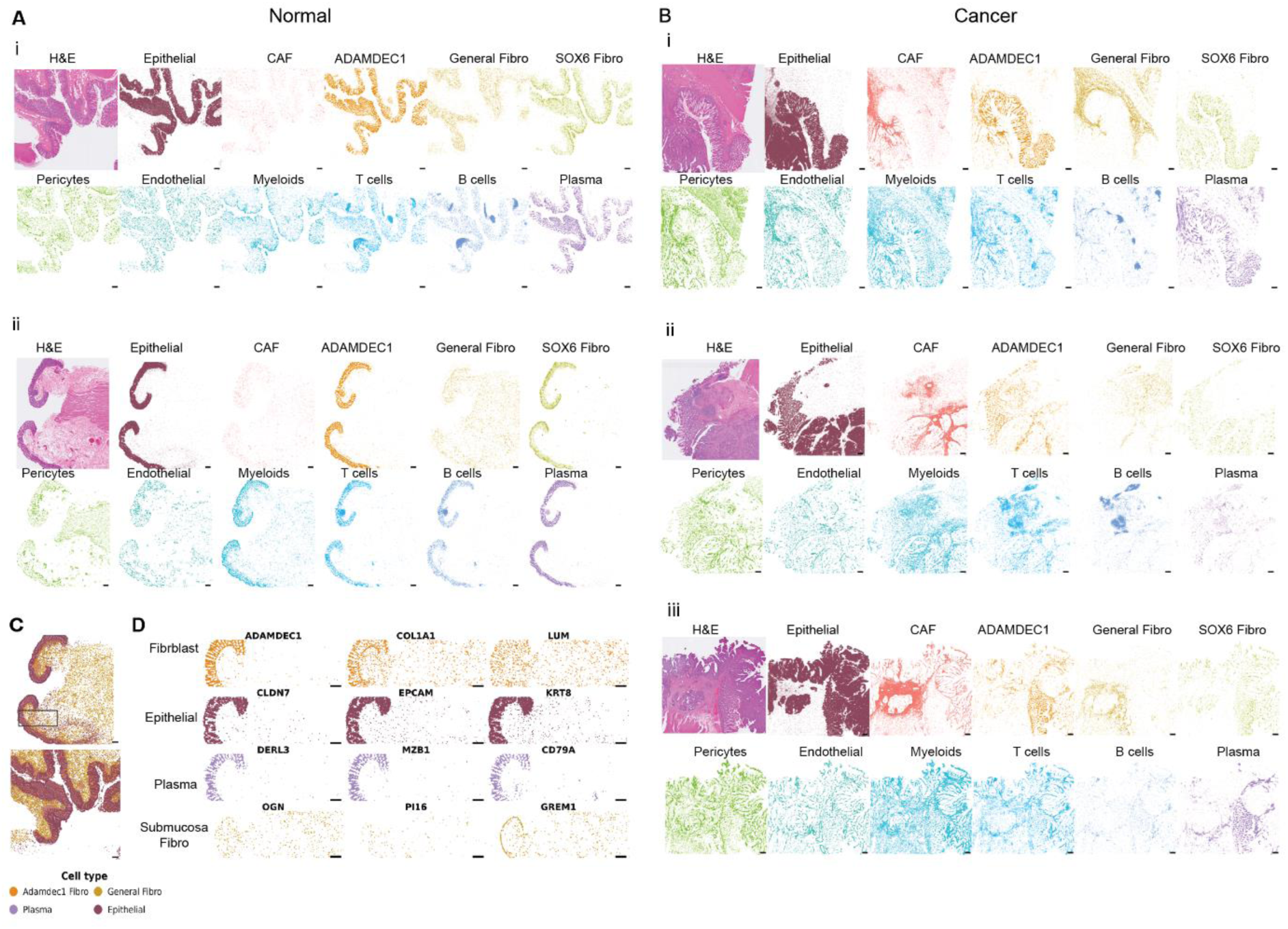
U-method markers define coherent spatial structures in VisiumHD sections. (A-B) Spatial projection of U-method derived cluster signatures in samples from CRC patients (n= 2 normal adjacent and n=3 tumor samples). Expression values are shown as raw averages of the top five U-markers per cluster, with spot alpha values ranging from 0.9 (any expression) to 1.0 (maximal expression). (C) Overlay of epithelial, plasma, ADAMDEC1^+^ fibroblast and General fibroblast signatures in 2 normal adjacent colon samples from CRC patients. Scale bar - 300 µm. (D) Comparison of the spatial projections of U-markers, canonical markers and general markers for epithelial, plasma, ADAMDEC1^+^ fibroblast and General fibroblasts. Scale bar - 200 µm.

To test the utility of the U-method across spatial transcriptomics datasets, we applied the full U-method analysis pipeline on scRNA-seq and Visium HD data from lung cancer. We processed publicly available scRNA-seq data from 22 lung adenocarcinoma patients^3^ using a standard Seurat workflow, and applied standard U-method parameters, similar to those used for the CRC, breast and PDAC datasets. The U-method identified UEGs for each subpopulation (Supplementary Figure 3A-D), recovering both canonical lineage markers and additional high-scoring cluster-specific candidates. We then projected the resulting UEG signatures (top 5 UEGs, U-score > 0.2) onto a publicly available lung adenocarcinoma Visium HD section (https://www.10xgenomics.com/datasets/visium-hd-cytassist-gene-expression-human-lung-cancer-post-xenium-expt). This analysis showed coherent spatial patterns (Supplementary Figure 3E). myCAFs were the dominant fibroblast population within the tumor stroma, whereas iCAFs and normal fibroblasts localized to surrounding non-tumor regions. Muscle signatures outlined both the thick structural wall of the lung and vascular scaffolds. Plasma cells and T/NK cells were dispersed along immune routes, while T cells and B cells co-localized in niches reminiscent of antigen-presenting or lymphoid-like sites. Beyond these cellular compartments, the U-method captured higher-order organization at the airway level. Monocytes localized within the airway lumen, directly bordered by ciliated epithelial cells and encased by smooth muscle. This layered arrangement recapitulates the architecture of immune-epithelial-muscle crosstalk that regulates airway biology^26^. Importantly, this organization was not imposed by manual tuning or annotation but instead emerged directly from probability-based marker scoring and projection. While this spatial analysis was performed on only one patient section, it demonstrates the utility of the U-method in revealing biologically meaningful structures.

A key practical feature of the U-method is that it can operate on any biologically meaningful partitioning of the data, without requiring that clusters represent fully resolved or homogeneous cell states. To illustrate this flexibility, we relabeled the epithelial/cancer compartment in the CRC dataset^2^ which was initially clustered as a single “Epithelial” group without subdivision by sample origin. Rather than reclustering epithelial cells, we used clinical metadata to relabel cells derived from tumor samples as “Cancer” and those from adjacent normal mucosa as “Epithelial.” This resulted in one relatively pure epithelial group and a cancer group which in both transformed and non-transformed epithelial cells coexist (Supplementary Figure 4A). We intentionally did not further subdivide the cancer group, allowing the analysis to test whether probability-based scoring could capture consistent expression differences under this coarse partitioning. Under this labeling, the U-method identified genes whose expression probabilities consistently distinguished cancer from epithelial cells despite substantial overlap in UMAP space (Supplementary Figure 4A). In epithelial cells, CA2 and VSIG2 received the highest U-scores, consistent with their known association with normal colon epithelium^5,7^. In the cancer-labeled group, S100P and LCN2 scored highest (Supplementary Figure 4B-C), in line with their reported upregulation in colorectal cancer^27–29^. When projected onto Visium HD colorectal sections, these marker sets distinguished the normal epithelial cells from adjacent cancer cells, illustrating that probability-based marker scoring remains effective even when applied to coarse or metadata-defined partitions (Supplementary Figure 4D).

### U-method driven spatial analysis reveals coordinated cell-type localization in normal and tumor tissues

The spatial maps guided by the U-method in colon and lung tissue revealed interpretable relationships at tissue scale. In normal colon mucosa, epithelial cells, plasma cells, and a distinct ADAMDEC1-positive fibroblast population consistently appeared in close proximity, forming a structured mucosal layer, a pattern that was reproducible across samples (Figure 3A and 3C). In tumor sections, signals were more intermixed and less distinctly layered, consistent with disrupted organization (Figure 3B). To quantify the observed spatial changes and interactions between stromal cells and epithelial structures, we combined Loupe Browser K-means clustering (https://www.10xgenomics.com/support/software/loupe-browser/latest) with U-method marker signatures, applying a radius-based enrichment approach (Supplementary Figure 5A). Using the epithelium-dominated Loupe cluster as the reference region, we defined all spots located within 200 pixels of the nearest epithelial spot as the IN group, and all other spots were considered the OUT group.

For each biological cluster defined by its U-method marker set, we first identified pure spots, defined as those expressing at least one marker of that cluster and none of the markers associated with other clusters. We then quantified the proportion of these pure spots IN versus OUT of the radius-defined region. When applied across all samples, this analysis produced stable and interpretable spatial patterns (illustrated for matched normal and tumor tissue from the same patient; Supplementary Figure 5B-C). Relative risk (RR) values were then calculated to compare the likelihood of observing each pure cluster inside versus outside the defined region (Supplementary Figure 5D).

Epithelial spots were strongly enriched inside the radius region, as expected, confirming that the RR framework can reliably quantify spatial enrichment based on U-method marker expression (Figure 3A, Supplementary Figure 5D). Several stromal populations also displayed distinct spatial associations with epithelial structures. In normal colon, ADAMDEC1^+^ fibroblasts and SOX6^+^ fibroblasts were consistently more abundant inside epithelial proximal regions, whereas general fibroblasts were more common outside these regions. Plasma cells were also enriched inside epithelial regions whereas mature B cells did not show any enrichment. These spatial characteristics were massively altered in cancer. The enrichment of ADAMDEC1^+^ fibroblasts and plasma cells to epithelial regions was dramatically reduced. In fact, the enrichment of most stromal and immune cell types increased in the OUT regions in cancer compared to normal adjacent tissue, most likely reflecting the reorganization of the tumor microenvironment to support changes in epithelial structure during tumor development captured by this radius-based analysis. Together, these analyses demonstrate how U-method derived markers enable quantitative mapping of spatial relationships between cell types, revealing coordinated tissue organization and its disruption in cancer.

## Discussion

In this study, we introduce the U-method, a probability-based framework for identifying uniquely expressed genes (UEGs) within any pre-analyzed scRNA-seq dataset. By contrasting a gene’s detection probability within a cluster with its highest probability in any alternative cluster, the method prioritizes detection consistency over expression magnitude. Applied across colorectal, breast, pancreatic, and lung cancer datasets, the U-method consistently recovered canonical lineage markers and produced concise marker sets that were reproducible across independent cohorts and clustering resolutions. These markers were directly suitable for spatial projection without smoothing or model-based inference. This method is fast, transparent and robust, making it a practical starting point for marker discovery and spatial validation.

A central challenge in single cell transcriptomic analysis is distinguishing robust biological signal from noise in sparse, high-dimensional data. Although differential expression methods are powerful for detecting quantitative differences, magnitude-based statistics can be sensitive to sequencing depth, technical noise, normalization choices, and the influence of a small number of high-count cells. In practice, cluster identity is often defined not by an exhaustive catalogue of expression changes, but by a limited set of markers that behave consistently across cells and datasets. These markers provide a stable reference for cell type annotation: once established, they enable downstream analyses to proceed without continual reassessment of the full expression landscape. Detection probability directly captures this reproducibility at the single-cell level. By using a maximum-outside contrast, the U-method emphasizes genes that exhibit polarized, cluster-specific detection patterns while filtering out marginal effects that depend on modeling choices or subtle shifts in mean expression. The resulting marker sets are deliberately conservative, yet stable and interpretable across analytical contexts.

Across tumor types, probability-based scoring yielded stable marker rankings. Core canonical genes such as AIF1, MZB1, MS4A1, VWF, RGS5, CD3D, and CD3E consistently appeared among the highest-scoring genes, often in similar rank order across tissues. In addition, the U-method identified biologically plausible markers that are frequently overlooked by magnitude-based methods, including genes with modest but highly specific expression patterns, such as DERL3 in plasma cells. This cross-dataset reproducibility was observed without dataset-specific tuning and across independently generated cohorts. Rather than maximizing the total number of detected differences, the U-method emphasizes markers that persist across datasets, thereby supporting consistent cell-type annotation and reducing dependence on study-specific effects.

Projection of UEGs onto Visium HD sections confirmed the biological relevance of the selected markers. Using only raw average expression of top UEGs, without smoothing or deconvolution, spatial maps recapitulated coherent epithelial, stromal, and immune compartments in normal tissue, and demonstrated how these patterns are disrupted in tumor sections. In the lung, the U-method captured both cellular organization and higher order airway architecture. These projections were generated directly from single-cell-defined signatures, without tuning to spatial data, supporting the use of UEGs for hypothesis generation and tissue-level interpretation.

The U-method is complementary to, rather than competitive with, high-resolution differential expression analyses. Magnitude-based approaches remain essential for pathway enrichment, mechanistic interpretation, and modeling of continuous trajectories or gradient-like expression changes. However, such analyses benefit from a stable set of identity-defining markers established in advance. By emphasizing detection consistency, the U-method provides this foundation while remaining agnostic to downstream statistical frameworks.

The U-method performs best when clustering reflects meaningful biological structure. If clusters are under-resolved, distinct states may be merged, reducing marker specificity. If over-resolved, coherent populations may split, diluting signals. In practice, we found that a global U-score threshold of 0.2 preserved subtle but reproducible contrasts, such as those between related epithelial states, while maintaining cross-dataset stability. Higher thresholds restrict selection to only the most exclusive markers, whereas lower thresholds increase sensitivity at the expense of specificity. Notably, U-score distributions exhibit a useful diagnostic property: coherent populations generate sharp, right-skewed profiles, while mixed or heterogeneous groups produce flatter distributions. This behavior suggests that detection consistency may itself inform assessments of cluster coherence. While not exploited directly in this study, this U-method feature could be formalized as a clustering quality metric.

The probability-based formulation also admits natural extensions. Replacing the maximum-outside term with a minimum-outside probability enables identification of cluster-specific absence of detection. Although this extension is not implemented in the current package and was not evaluated here, it follows naturally from the method’s structure and may be useful when the absence of a gene distinguishes closely related cell states. Spatial projection of uniquely unexpressed signatures is less intuitive, but negative-expression maps can be generated when biologically meaningful.

More broadly, the U-method formalizes detection consistency as a complementary dimension of marker specificity. Expression magnitude and detection probability capture distinct statistical properties of gene behavior: magnitude reflects quantitative differences in expression levels, whereas detection probability reflects stability of presence across individual cells. The choice between them reflects a bias-variance tradeoff in marker selection. By prioritizing detection consistency, the U-method sacrifices sensitivity to small quantitative shifts in favor of robustness and cross-dataset reproducibility. This perspective reframes marker discovery as the identification of features that are stable enough to anchor biological interpretation rather than exhaustive enumeration of differential effects.

In summary, the U-method provides a simple, interpretable, and robust approach for marker discovery across single-cell and spatial transcriptomics datasets. By relying on detection probability, it avoids assumptions tied to expression magnitude and normalization, and it consistently yields markers that align with biological structure and spatial organization. Its speed and conceptual clarity make it well suited for exploratory analysis, structured validation, and integration into diverse single-cell and spatial transcriptomics workflows.

## Methods

### Overview of the U-method

The U-method is a probability-based framework for identifying uniquely expressed genes (UEGs) within any clustering of scRNA-seq data. For each gene and each cluster, the method compares the detection probability in that cluster with the highest detection probability observed in any other cluster. This maximum-outside comparison is central to the approach, since it asks whether a gene is more frequently expressed in the target cluster than in its strongest competitor, rather than contrasting a cluster with the average behavior of all other populations.

This formulation reflects the binary nature of single-cell measurements, where many biologically relevant genes are effectively on in one population and off in others. It avoids assumptions about expression magnitude or distribution, and it is independent of the specific clustering algorithm and normalization strategy. When high-resolution spatial data are available, U-method markers can be projected onto Visium HD tissue sections by averaging raw expression of top UEGs per cluster, leveraging the fact that a cell-type-specific signature identified in single-cell data can be approximated spatially using unscaled average expression. All analyses were performed using the R package U-method, which provides reference implementations of the core steps.

### Expression probability scoring (FindUniqueMarkers)

#### Data inputs

Let X be a gene-by-cell matrix of raw or normalized counts, and let each cell have an assigned cluster label C ∈ {1,…,K}. The U-method operates on any precomputed clustering and does not constrain upstream preprocessing.

#### Expression probability

For every gene g and cluster c, we define the in-cluster probability of expression as

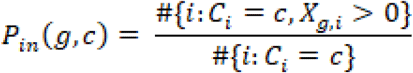

To evaluate how often gene g is expressed outside cluster c, we compute its maximum probability in any other cluster,

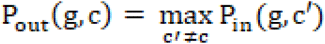

Thus, each gene–cluster pair is contrasted against the strongest competing cluster, not against an average of all others. This “worst competitor” formulation ensures that a gene is considered specific only if its detection probability in the target cluster exceeds that of every alternative cluster.

#### U-score

The U-score quantifies the cluster specificity of gene g in cluster c,

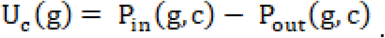

The U-score ranges from −1 to 1. A value near 1 indicates that the gene is frequently detected in cluster c and rarely or never detected in any other cluster. Scores near 0 indicate low specificity, and negative values reflect higher detection probabilities in other clusters. In practice, U-scores provide the primary basis for identifying and ranking UEG candidates and do not require expression scaling or modeling of continuous expression levels.

### Standardization and statistical evaluation of U-scores

Although the U-score itself is the core specificity measure, we include an optional hypothesis-testing layer for use in cases where strict exclusivity is required, such as selecting a single marker for immunofluorescence staining.

#### Standardization

For each cluster c, U-scores across all genes are standardized to obtain a cluster-specific z-score distribution,

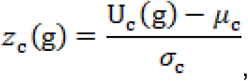

where μ_c_ and σ_c_ are the mean and standard deviation of U_c_(g) over all genes in cluster c. This standardization facilitates an approximate null model while preserving the ordering induced by U-scores.

#### Statistical significance

One-sided p-values are computed by comparing z_c_(g) to the standard normal distribution, and Benjamini–Hochberg correction is applied within each cluster. Under this approximation, a statistically significant gene is interpreted as one whose probability of expression in cluster c is conclusively higher than in its strongest competitor cluster. This is a deliberately strict criterion and is intended for applications that require markers that are as exclusive as possible.

In practice, the empirical distribution of U-scores includes many non-specific or outwardly enriched genes, which generates a lower tail with inflated variance relative to the ideal null, in which expression probabilities are identical across clusters. As a result, the normal approximation yields conservative p-values. Correcting this bias exactly would require computationally intensive procedures such as permutation-based null estimation or explicit modeling of the null gene class, which is at odds with the goal of a fast, hands-on method.

For broader marker discovery, we therefore rely on the magnitude of the U-score rather than on p-values. In this study, genes with U_c_(g) >0.2 were considered robustly enriched and identified as uniquely expressed genes, providing a practical heuristic that captures genes with consistent probability-based specificity without requiring slow null-model estimation. Users who seek the most strictly unique markers can use the significance filter or use alternative methods to estimate the U-scores p-values under a more realistic null hypothesis, while users who need richer marker sets can rely on U-scores and thresholds alone.

### Integration with Visium HD spatial transcriptomics data (CreateImageData)

To visualize U-method markers in spatial context, we converted Visium HD outputs into spatial Seurat objects. Filtered gene count matrices were loaded using Read10X, and spatial coordinates and spot metadata were obtained from the tissue_positions.parquet files. The CreateImageData function merges these components into a Seurat object that contains raw expression values for the selected markers alongside their spatial coordinates.

Optionally, spots can be annotated with binary marker expression (expressed or not expressed) based on non-zero counts. This representation supports both continuous and presence/absence–based analyses while keeping the spatial structure explicit.

### Signature scoring and spatial projection (UmethodSignatureMap)

For each cluster in the scRNA-seq reference, we defined a cluster-level signature as the set of its top UEGs, typically the top five genes ranked by U-score, or all available markers when fewer were present. Spatial projection of these signatures onto Visium HD sections proceeds in three steps.

First, for each Visium HD spot j and cluster c, we compute a raw expression score

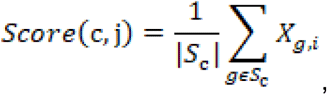

where Sc is the set of marker genes for cluster c, and □□, □ is the raw count of gene g at spot j. No smoothing, scaling, deconvolution, or normalization is applied at this stage.

Second, each spot is assigned to the cluster whose signature score is maximal,

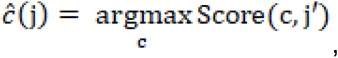

yielding a winner-takes-all spatial classification map.

Third, these scores and assignments can be used to generate spatial plots that display either the average expression of cluster signatures or a winner-takes-all spatial classification. In this study, we focus on visualizing the raw average expression of each cluster signature across Visium HD sections, rather than the per-spot classification labels. This choice highlights the underlying expression patterns of the U-method markers and allows direct inspection of their spatial coherence without imposing a hard assignment. For applications that require displaying all clusters simultaneously, such as interactive plotly maps that benefit from a discrete categorical layer, the classified version provides a practical alternative.

The UmethodSignatureMap function implements this procedure and outputs both per-spot classification labels and long-format tables of per-spot, per-signature scores.

#### Spatial visualization

Spatial maps were generated using ggplot2, with each spot plotted in its physical coordinates. Spots were colored either by their assigned cluster label or by raw average expression of a given marker set. Alpha values were used to emphasize regions with higher expression, with alpha ranging from 0.9 for any detectable expression to 1.0 for maximal expression. Only non-zero expression values were plotted. No model fitting, kernel smoothing, or dimensionality reduction was applied during spatial visualization, allowing a direct view of how U-method signatures map onto tissue space.

### Radius-based spatial enrichment analysis

To quantify the spatial localization of U-method–defined cell states relative to epithelial regions, we combined Loupe Browser K-means clustering with marker-based spot annotation.

For each Visium HD section, we selected the Loupe K-means cluster enriched for epithelial features as a reference region. For every spot, we computed the Euclidean distance to the nearest epithelial spot. Spots within 200 pixels of the reference were labeled in-radius, and all other tissue spots were labeled out-radius.

For each biological cluster defined by its U-method marker set, we identified pure spots as those expressing at least one marker of that cluster and none of the markers of other clusters, applying a simple negative-selection gating step to isolate the corresponding cell state. For each cluster, we then constructed a 2×2 contingency table comparing in-radius and out-radius occupancy by these gated pure spots and computed a relative risk statistic. For cross-sample comparison, the log RR values themselves were used to generate boxplots summarizing enrichment patterns across normal and tumor tissues.

### Cross-dataset marker reproducibility analysis

To assess marker reproducibility across independent datasets, we applied the U-method and a Wilcoxon-based differential expression approach to four publicly available scRNA-seq datasets, two from breast cancer^5,7^ and two from PDAC^4,6^. All datasets were processed using Seurat with standard normalization, variable feature selection, dimensionality reduction, and clustering, with integration where appropriate.

To enable direct comparison with the colorectal analysis, we applied the same stromal subclustering strategy across datasets. Fibroblasts and pericytes were first identified based on canonical markers derived from U-method analysis of the full dataset at low clustering resolution. These cells were then subsetted and reclustered independently using the same Seurat workflow applied to the colorectal stroma. The resulting stromal subclusters were used to refine the original low-resolution fibroblast and pericyte populations in the full dataset. Mixed clusters, such as “apCAF” and “Stromal Else”, were excluded from marker analysis to avoid dilution of cluster-specific signals, while remaining included in visualizations to highlight their heterogeneous expression profiles and justify their exclusion.

For the U-method, we selected genes with U-score greater than 0.2 and retained up to the top five UEGs per cluster. For the Wilcoxon analysis, we used Seurat’s FindAllMarkers with only.pos = TRUE and default options, retaining genes with adjusted p-value lower than 0.05 and absolute log2 fold change greater than 2 and retained up to the top five DEGs per cluster. Within each cluster, markers were ranked by U-score (U-method) or log fold change (Wilcoxon), and the top ten markers were used to compute pairwise overlap assessment across matched cluster types between datasets. Heatmaps of these indices were used to visualize marker overlap and reproducibility.

### Software implementation and availability

The U-method is implemented in the R package Umethod, which provides three core functions:

● FindUniqueMarkers, which computes expression probabilities, U-scores, z-scores, and optional p-values for each gene–cluster pair using the maximum-outside contrast.
● CreateImageData, which constructs spatial Seurat objects by integrating Visium HD count matrices with spatial metadata and marker expression.
● UmethodSignatureMap, which computes spatial signature scores, performs winner-takes-all spot classification, and outputs tables suitable for visualization and downstream analysis.

The package and analysis pipeline scripts are available at: https://github.com/YanuvS-Dev/Umethod.

## Data Availability

All datasets generated and analyzed in this study are publicly available.

The raw scRNA-seq data for CRC data are available in the European Genome-phenome Archive database (EGAS00001003779, EGAS00001003769). The breast cancer RNA sequencing data are available with accession number E-MTAB-10607 on the ArrayExpress database and in the European Genome-phenome Archive database EGAS00001005173, while PDAC RNA sequencing data are available in on GEO with accession number GSE205013 and Genome Sequence Archive under project PRJCA001063.

The Visium HD colorectal cancer samples (CRC1, CRC2, CRC5) and the corresponding normal tissues (NAT3, NAT5) were obtained from the 10x Genomics Human Colon HD Dataset and the Visium HD Human Colon Cancer Dataset. Raw spatial transcriptomics data, including count matrices and spatial coordinates for the 8 µm resolution captures (https://www.10xgenomics.com/platforms/visium/product-family/dataset-human-crc).

Marker tables and U-method results, including cluster-level unique expression gene (UEG) signatures, are available in Supplementary Table 1. Loupe Browser–derived K-means classifications used in the spatial enrichment analysis (KM2CRC1, KM2CRC2, KM2CRC5, KM3NAT3, KM2NAT5) are provided with this manuscript as Supplementary Table 2. Radius-based spatial enrichment analysis RR outputs are provided with this manuscript as Supplementary Table 3.

Additional scripts and analysis workflows are available upon request.

## Code Availability

The U-method R package is openly available at: https://github.com/YanuvS-Dev/Umethod

Analysis pipeline code is available at: https://github.com/YanuvS-Dev/Umethod.

## Supporting information

U-method marker statistics across scRNA-seq datasets.

Loupe Browser K-means spatial annotations.

Radius-based spatial enrichment analysis results.

---

**Supplementary Figure 1.**
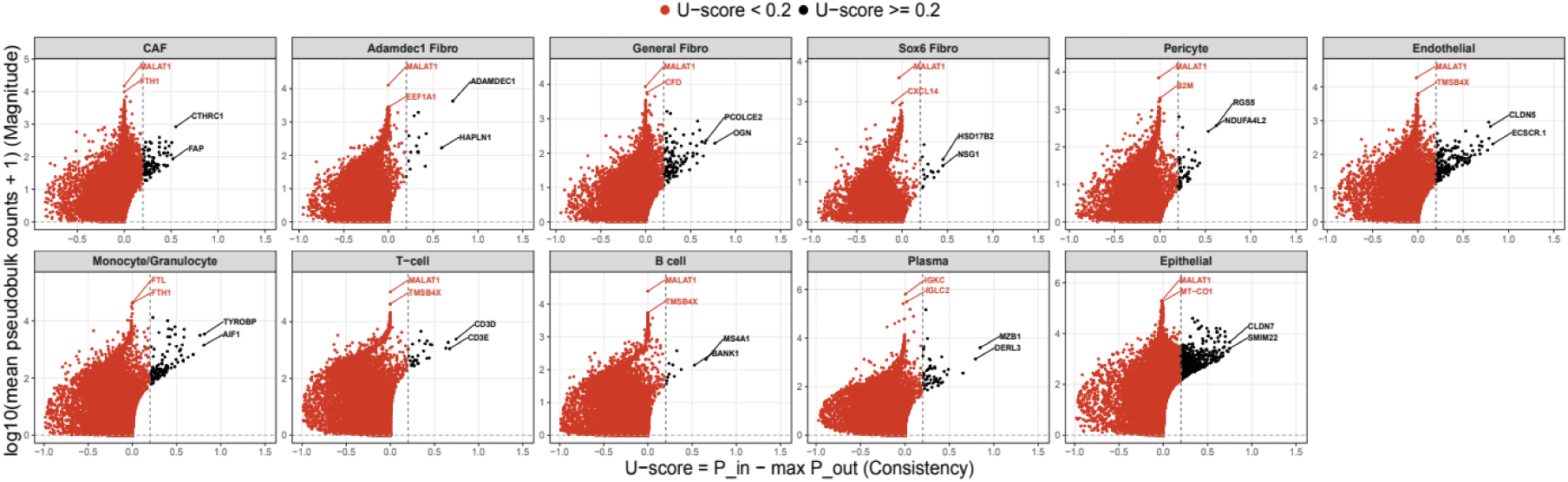
U-scores and pseudobulk aggregation-based markers measure different features in colorectal cancer scRNA-seq data. Scatter plot showing U-scores (X-axis) and log 2 mean pseudobulk counts for clusters identified in the CRC dataset ^2^(Y-axis; For each gene, expression counts were aggregated within each sample and cluster, transformed as sum log(x + 1), and averaged across samples to obtain a cluster-level pseudobulk expression summary). Each point represents a gene. Genes with U-score greater than 0.2 are highlighted.

**Supplementary Figure 2.**
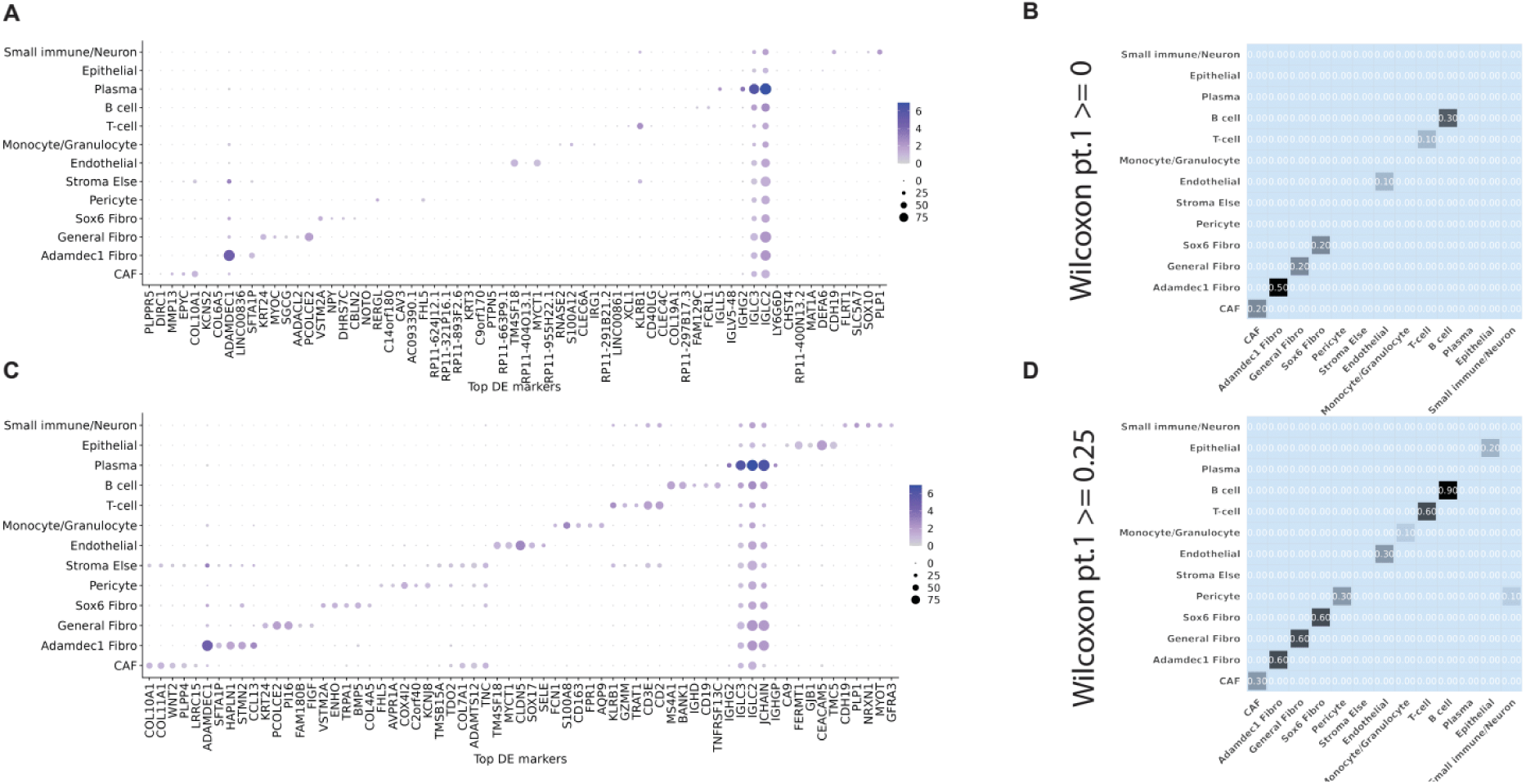
Low detection frequency explains discrepancies between Wilcoxon-derived and U-method markers. (A) Wilcoxon results with standard thresholds (|log2FC| > 2, adjusted p < 0.05), on the CRC data^2^ with stromal subclusters. (B) overlap matrix of Wilcoxon top 10 markers and U-method top 10 markers (C) Wilcoxon results with additional filtering (pct.1 > 0.25). (D) overlap matrix of Wilcoxon top 10 markers with additional filtering and U-method top 10 unfiltered markers.

**Supplementary Figure 3.**
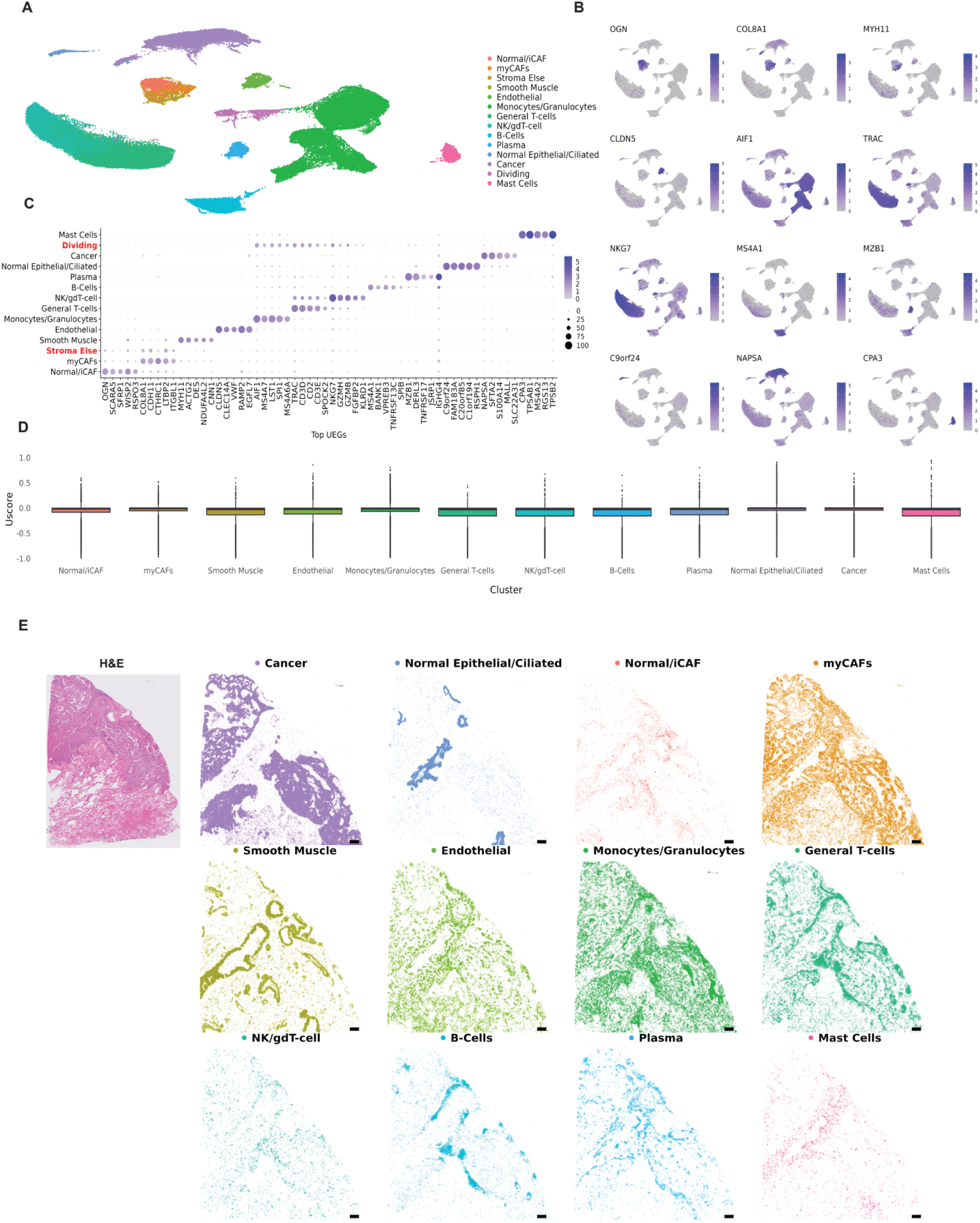
The U-method reveals robust markers and spatial organization in lung adenocarcinoma data. (A) U-method analysis of scRNA-seq data from lung adenocarcinoma^3^ annotated by cell type. UMAP projection of all cell clusters and fibroblast and T-cell subclusters is shown (B) Feature plots showing the top U-markers of the main cell clusters shown in A. (C). Unscaled Dot plots of the top uniquely expressed genes per cluster. (D) Boxplots of U-score distributions across clusters and subclusters. (E) Projection of the average expression of identified markers onto a VisiumHD lung adenocarcinoma sample (https://www.10xgenomics.com/datasets/visium-hd-cytassist-gene-expression-human-lung-cancer-post-xenium-expt).

**Supplementary Figure 4.**
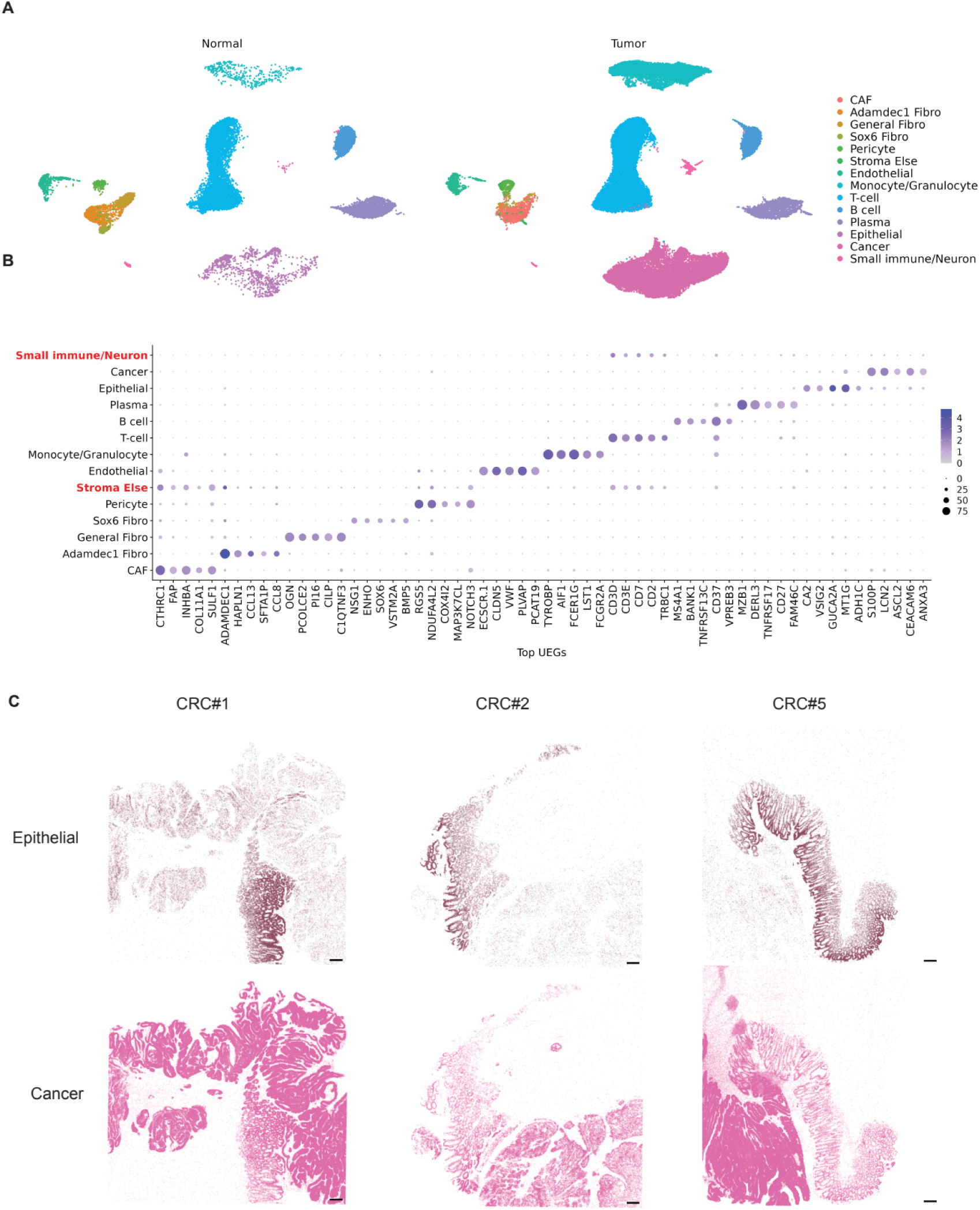
Projection of scRNA-seq-derived UEG average expression separates epithelial and cancer states in spatial transcriptomics. (A) UMAP projections of scRNA-seq data from human colorectal cancer (CRC) and adjacent normal colon tissue^2^ split by sample origin, normal or tumor, analyzed by Seurat and annotated by major cell type. (B) Unscaled Dot plot of the top uniquely expressed genes per cluster. (C) Spatial maps showing the projection of uniquely expressed gene (UEG) signatures derived from scRNA-seq of epithelial and cancer cell clusters onto Visium HD CRC tissue sections. For each spot, signature values were computed as the raw average expression of U-method-identified marker genes associated with the epithelial or cancer cluster. No smoothing, normalization, or deconvolution were applied. Spots are plotted at their physical coordinates, with color intensity reflecting the signature value and transparency indicating detectable expression. Scale bar - 300 µm.

**Supplementary Figure 5.**
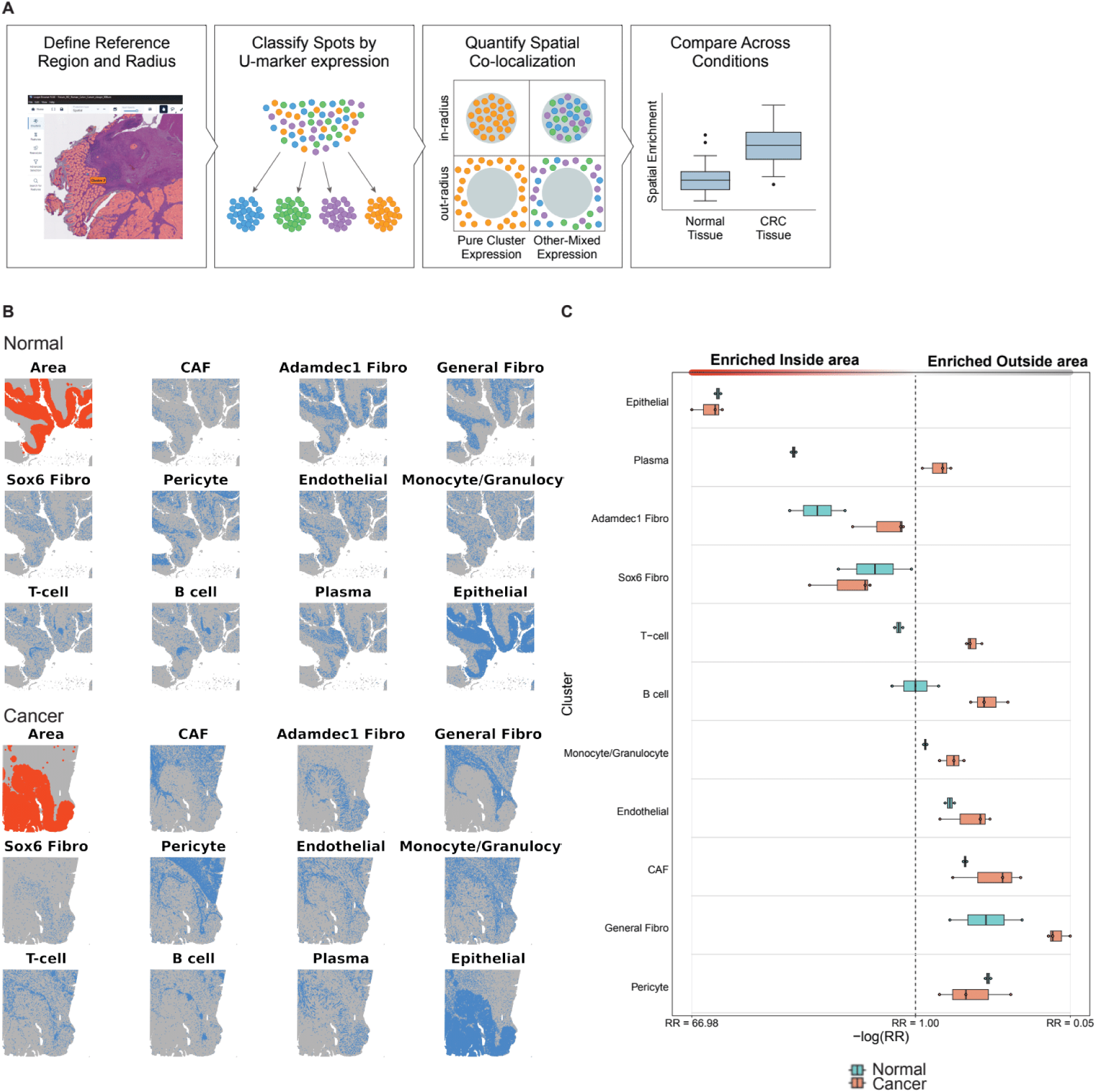
The U-method enables quantitative analysis of spatial relationships between cell types. (A) Schematic representation of the computational pipeline. (B-C) Representative spatial maps of matched normal and tumor samples from the same patient, showing IN-radius epithelial regions (orange) and pure cluster-expressing spots (blue). (D) Relative risk values across clusters and samples, reflecting enrichment of specific cell types near epithelial regions.

